# Space-independent community and hub structure of functional brain networks

**DOI:** 10.1101/590935

**Authors:** Farnaz Zamani Esfahlani, Maxwell A. Bertolero, Danielle S. Bassett, Richard F. Betzel

**Affiliations:** Department of Psychological and Brain Sciences, Indiana University, Bloomington, IN 47405; Department of Bioengineering, University of Pennsylvania, Philadelphia, PA 19141; Department of Electrical & Systems Engineering, University of Pennsylvania, Philadelphia, PA 19141; Department of Neurology, University of Pennsylvania, Philadelphia, PA 19141; Department of Psychiatry, University of Pennsylvania, Philadelphia, PA 19141; Department of Physics & Astronomy, University of Pennsylvania, Philadelphia, PA 19141; Cognitive Science Program, Indiana University, Bloomington, IN 47405; Program in Neuroscience, Indiana University, Bloomington, IN 47405; Network Science Institute, Indiana University, Bloomington, IN 47405

## Abstract

Coordinated brain activity reflects underlying cognitive processes and can be modeled as a network of inter-regional functional connections. The most costly connections in the network are long-distance correlations that, in the absence of underlying structural connections, are maintained by sustained energetic inputs. Here, we present a spatial modeling approach that amplifies contributions made by long-distance functional connections to whole-brain network architecture, while simultaneously suppressing contributions made by short-range connections. We use this method to characterize the long-distance architecture of functional networks and to identify aspects of community and hub structure that are driven by long-distance correlations and that, we argue, are of greater functional significance. We find that based only on patterns of long-distance connectivity, primary sensory cortices occupy increasingly central positions and appear more “hub-like”. Additionally, we show that the community structure of long-distance connections spans multiple topological levels and differs from the community structure detected in networks that include both short-range and long-distance connections. In summary, these findings highlight the complex relationship between the brain’s physical layout and its functional architecture. The results presented here inform future analyses of community structure and network hubs in health, across development, and in the case of neuropsychiatric disorders.

## INTRODUCTION

Cognitive and psychological processes are underpinned by the coordinated activity of spatially distributed brain areas [1]. This coordination pattern can be estimated from observed brain activity and quantified as the statistical dependence of brain regions’ activity on one another [2]. The set of all such measurements can be modeled as a functional network and analyzed using graph theoretic methods [3, 4].

Analysis of functional networks has revealed a number of key features of brain organization. Among the most salient are the brain’s modular structure [5, 6] and the presence of hub regions – brain areas whose connections span modular boundaries [7, 8]. Both of these features can be interpreted in the context of brain function: modules reflect units for performing specialized information processing while hubs reflect the integration of that information from one module to another [9, 10].

In general, the organization of functional networks (including hubs and modules) is guided and constrained by many factors. One important factor is the brain’s underlying anatomical network of white-matter fiber tracts [11–13]. This anatomical network plays an important role in shaping activity across the brain [14, 15] and is, itself, subject to strong metabolic and spatial constraints [16–18]. These constraints effectively place soft limits on the possible number, length, and volume of structural connections [19].

The mapping of structural connectivity (SC) to functional connectivity (FC) is complex [13, 20–22] and constraints on wiring-cost propagate to the level of FC, leading to gradients of distance-dependent inter-areal correlations wherein activity recorded from nearby brain areas tends to be more strongly correlated than that of distant areas [6, 23].

How do we disambiguate patterns of FC that arise out of functional necessity from those that arise as a consequence of space? One possibility is to emphasize long-distance FC while discounting short-range FC. Correlations between distant brain areas have, in general, little underlying structural support [11, 13, 20, 24, 25] and are instead maintained *via* inter-areal communication along multi-step, polysynaptic pathways, requiring sustained energy input [26–29]. Despite the energetic cost, many brain areas exhibit correlated activity over long distances. This observation suggests that the cognitive processes supported by these communication patterns cannot be easily subsumed by short-range connections, despite the fact that those short-range patterns would likely have stronger structural underpinnings and be more energetically efficient.

The principal aim of this study was to better characterize long-distance functional connectivity and to understand its contribution to the community structure and hub distribution of functional networks. To accomplish this goal, we developed a modeling framework that enabled us to generate synthetic regional time series whose correlation structure exhibited a prescribed distance-dependence [30]. While these synthetic networks exhibited typical patterns of short-range connectivity, they lacked the long-distance correlations that are also observed in functional networks. To shift focus onto those long-distance connections, we subtracted the elements of synthetic matrices from the corresponding elements in the observed functional connectivity matrix. The resulting network contained correlations among different brain regions that were stronger than expected, given the distance between those regions.

Surprisingly, we found that synthetic networks exhibited strong functional connections, both within and between well-characterized canonical brain systems, suggesting that short-range and long-distance connections make differential contributions to whole-brain functional connectivity. Next, we show that, after correcting for space, the participation coefficients of brain areas within primary sensory systems (somatomotor and visual) increase, suggesting that they may perform increasingly “hub-like” functional roles based on long-distance connectivity. We then use a data-driven approach for defining communities both with and without a distance correction. Overall, we find that the identified communities are similar across methods, with subtle differences in primary sensory systems but also higher-order cognitive systems, including salience and dorsal attention networks. In summary, these findings present a complex portrait of the relationship between space and functional connectivity. These results inform future studies of brain network communities and hubs, and may further clarify the role of functional connectivity in health, development, and disease.

## RESULTS

In this paper, we analyze the organization of long-distance functional connections, the results of which are presented in the following subsections. First, in the section **Modeling FC distance dependence**, we explain the basic features of our model and the procedure used to fit the model to observed data. In the next section, **Comparison with cognitive systems**, we introduce the distance-corrected functional connectivity matrix, focusing on where it differs from the observed, functional connectivity matrix. We also show that, given a set of canonical cognitive systems, distance-corrected functional connectivity forces us to rethink our current understanding of functional hub locations in cortex. Next, in the section **Incorporating distance-dependence into community detection tools**, we use the distance-corrected functional connectivity matrix as input to a community detection algorithm. With this algorithm, we detect communities in both distance-corrected and observed functional connectivity, and we compare the outputs. Finally, using detected communities, we describe variations in hub organization as a function of community size, reporting four distinct spatial patterns.

### Modeling FC distance dependence

In order to study long-distance functional connectivity, we first needed a method for generating synthetic functional connectivity with a prescribed distance dependence. In addition, we wished to ensure that the resulting synthetic functional connectivity matrix was admissible as a correlation matrix, being positive semidefinite and satisfying transitive relationships. Briefly, our strategy involved generating synthetic regional fMRI BOLD time series using phase-randomization procedures (Figure. 1a*i*). Because phase randomization was performed separately for every brain region, the resulting time series were uncorrelated, on average. To introduce spatial correlations, we defined a new time series for each region as a weighted sum of all other regional time series (Figure. 1a*ii*). Here, the weights were defined to be inversely proportional to the Euclidean distance between pairs of brain regions, so that nearby regions contributed more than distance regions. As a result, the magnitude of inter-regional correlations decreased monotonically as a function of distance (Figure. 1a*iii*). The rate of this decrease was modulated by a single parameter (Figure. 1b,c) and fit to observed data (Figure. 1d). The model is described in greater detail in **Materials and Methods**.

**FIG. 1.**
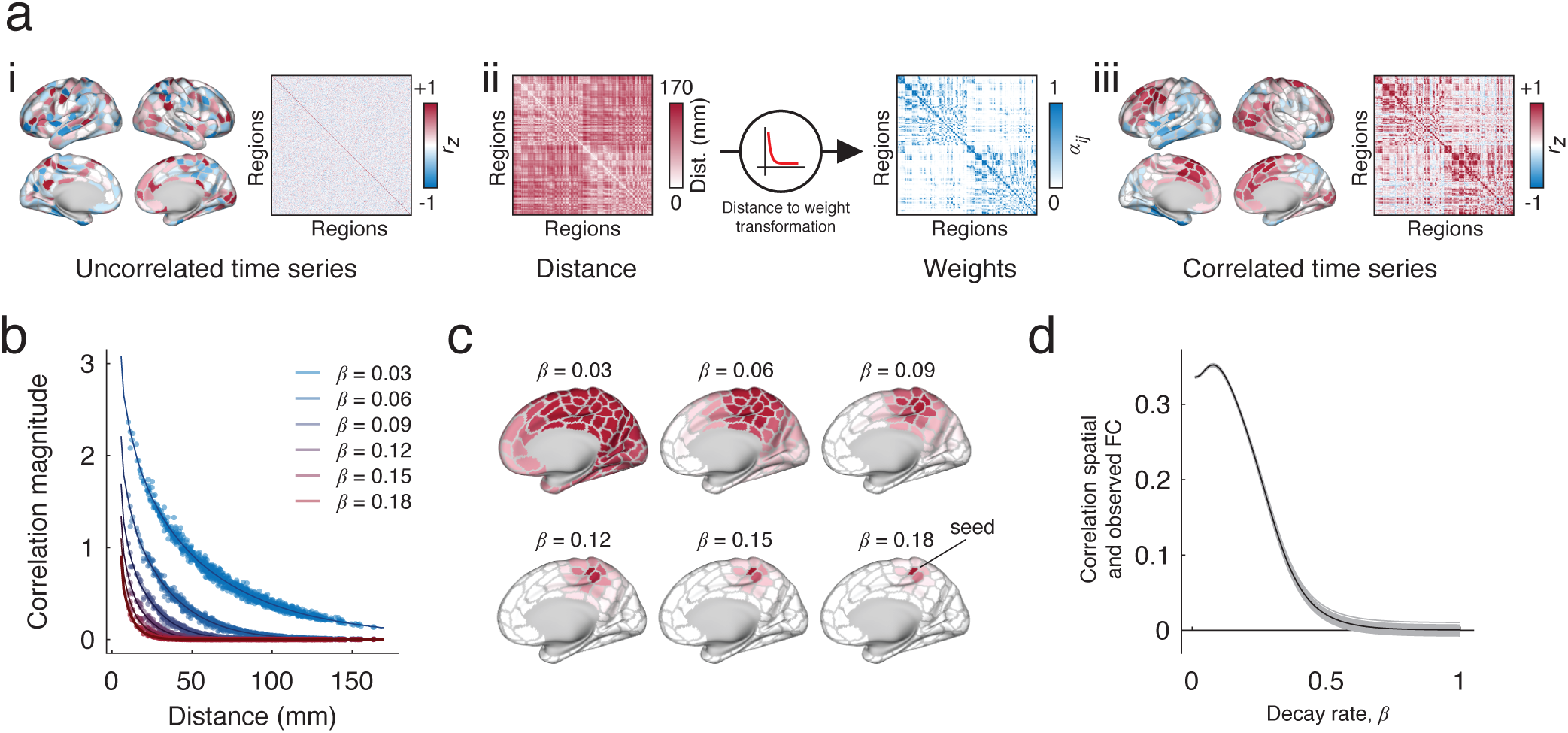
Procedure for constructing matrices with distance-dependent FC. (*a*) *i* : The procedure begins by introducing a random phase to the observed fMRI BOLD time series. This introduction is performed independently for each brain area, resulting in an uncorrelated surrogate time series. *ii* : Next, we generate a new surrogate time series for each brain area as the linear combination of all other time series. The weighted contribution of brain area *j*’s time series to that of area *i* depends on the distance between those two areas in three-dimensional Euclidean space, *D*_*ij*_. Specifically, this distance-to-weight transformation is modeled as a decaying exponential, 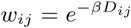, where *β* controls the decay rate. *iii* : This process results in spatially correlated time series. (*b*) The value of *β* controls the rate of the exponential decay and therefore parameterizes the extent to which time series exhibit more or less spatial correlation. For example, large values of *β* yield areal time series that are correlated with only their nearby neighbors; smaller values of *β* yield spatial correlations with much wider neighborhoods. (*c*) Examples of seed-based functional connectivity as we vary the value of *β*. (*d*) To select the value of *β*, we compare the similarity of the observed and spatially constrained FC patterns. The optimal value of *β* is the one that maximizes the correspondence between those matrices.

Our approach is similar, in spirit, to previous methods for generating distance-dependent correlation matrices using rational quadratic functions [31], which have proven useful in geospatial statistics [32]. Our approach, however, operates directly on time series whereas the approach of [31] and others is model-based. Both approaches generate positive-definite matrices with distance-dependent elements. Throughout this section, we use the synthetic matrices generated by this model as a sort of null condition that allows us to identify a set of spatially unexpected features [33].

### Comparison with cognitive systems

Functional connectivity can be used to map the brain’s organization at the level of systems [6, 34]. Analyses of this type generated cortex-wide maps in which brain areas or parcels are assigned to one or another system. These systems, in turn, are often interpreted in the context of brain and cognitive function; for example, some are thought to comprise collections of brain areas that support somatomotor function or for enacting cognitive control. Importantly, in conjunction with network measurements like participation coefficient, these systems have also been used to detect and classify hubs areas based on whether an area’s connections are distributed across many systems (hub) or concentrated within an area’s own system (non-hub) [7, 8, 35]. Here, we compared observed patterns of functional connectivity with those generated by the spatial model in order to gain insight into the contribution of long-distance connections to the brain’s system-level organization and hub structure [36].

The analyses in this section focused on three interdependent connectivity matrices: the observed pattern of functional connectivity, *FC*^*observed*^ (Fig. 2a); the synthetic patterns of connectivity generated by the spatial model, *FC*^*observed*^ (Fig. 2b); and the distance-corrected matrix of functional connections, *FC*^*corrected*^ (Fig. 2c), which was calculated as the element-wise difference of the observed and spatial matrices, *FC*^*observed*^ − *FC*^*spatial*^). As expected, when we reordered the *FC*^*observed*^ by system, we found that systems were cohesive with strong intra-system correlations (Fig. 2b). When we plotted *FC*^*spatial*^, we found that connections within many of the systems were much weaker in comparison (for example, the control networks Conta and Contb). On the other hand, we were surprised to find that a number of systems maintained their cohesiveness despite the fact that their connections reflect a spatial wiring rule rather than relevance to a cognitive or psychological process. The somatomotor and visual systems, for instance, as well as Contc, a sub-component of the control network, were dominated by strong short-range connectivity. In Fig. 2c, we show the effect of correcting for these distance effects, noting the now-attenuated connection weights within visual, motor, and control networks (see black arrows).

**FIG. 2.**
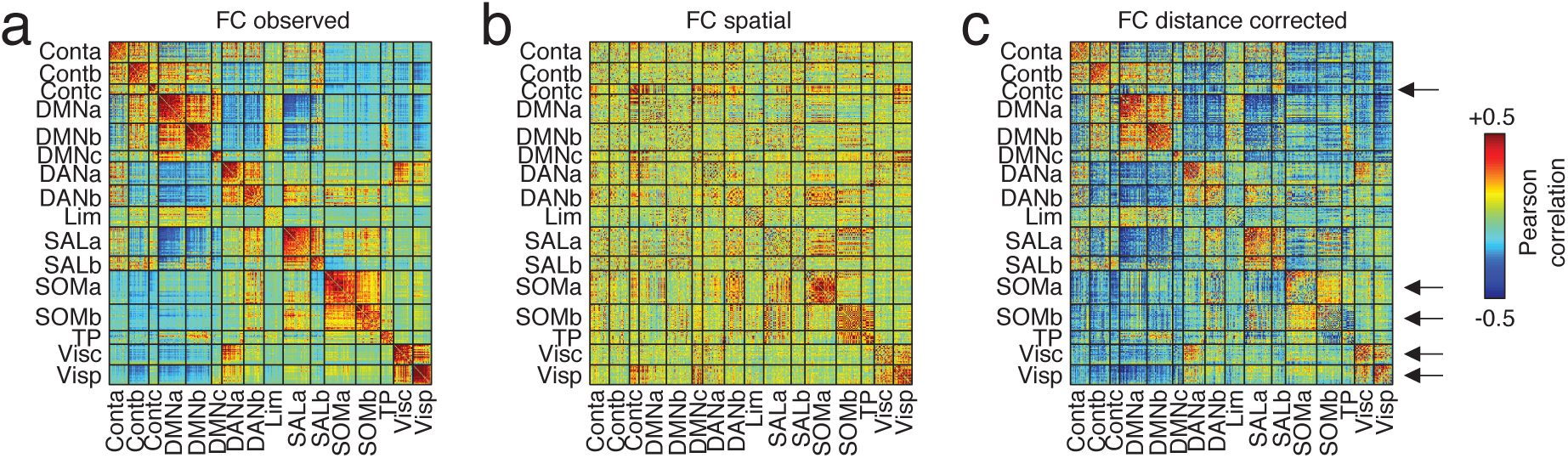
Differences in correlation structure. (*a*) The original (uncorrected) inter-regional correlation matrix ordered by cognitive system. (*b*) The correlation matrix obtained from the optimal spatial null model. (*c*) We correct for space-induced inter-regional correlation structure by subtracting the spatial null model matrix from the original matrix. Here, the arrows are used to draw attention to salient changes. The top arrow, for instance, shows that after correcting for distance, regions within the “Contc” system become weakly connected to each other. Similarly, in the original network, the correlation magnitude within somatomotor and visual systems are among the strongest across the brain. As with the control network component, these systems exhibit marked reductions in their internal connection strength following correction.

The observation of commonalities and differences between *FC*^*observed*^ and *FC*^*spatial*^ has implications for our understanding of brain function. In particular, it suggests that some network features may be driven differentially by short-range or long-distance connections. Here, we focus on the network property of “participation coefficient,” a local measure typically interpreted as an index of a region’s “hubness”. To identify areas whose observed participation coefficient may be driven predominantly by short-range connectivity, we computed each brain area’s signed participation coefficient [35] using the function `participation_coef_sign.m` in the Brain Connectivity Toolbox (https://sites.google.com/site/bctnet/) [4] with respect to the system assignments reported in [36] (Fig. 3a,b). We repeated this procedure for both the observed (Fig. 3c) and distance-corrected FC matrices. Next, to reduce bias from differences in average connection weight, we ranked participation coefficients and computed the difference in rank for each brain area (Fig. 3d,e). Interestingly, we find that the areas with the greatest increases in participation co-efficient are concentrated in visual and motor systems (*p <* 0.05; false discovery rate fixed at 5%), indicating that a greater proportion of connections from these areas now span system boundaries. These differences can be attributed to the fact that in the distance-corrected matrix, the weights of functional connections within somatomotor and visual networks are massively attenuated. That is, those connections are expected under the spatial null model. We also observed that the participation within components of the default mode, salience and ventral attention networks decreased significantly (*p <* 0.05; false discovery rate fixed at 5%).

**FIG. 3.**
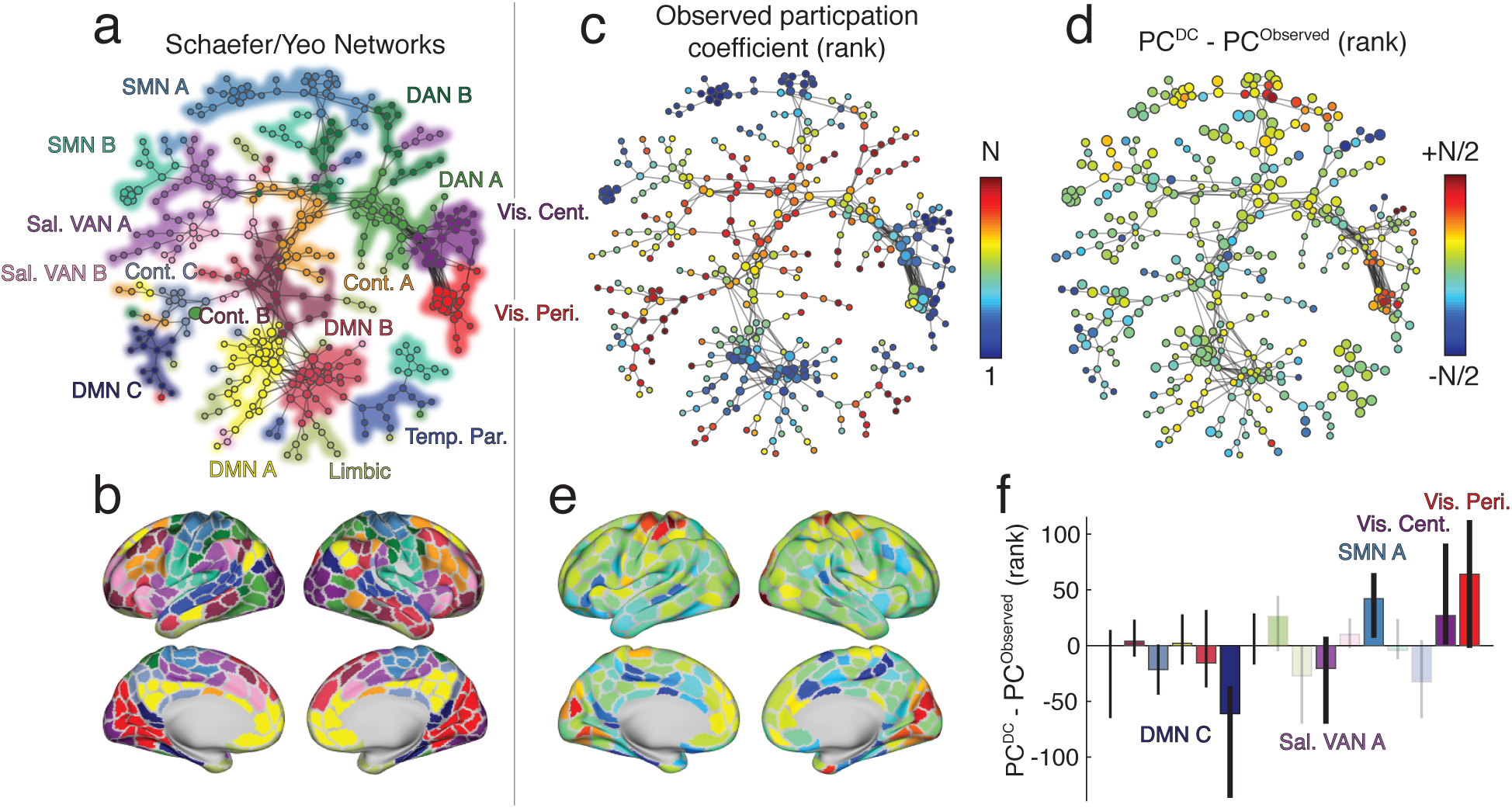
Effect of distance correction on canonical brain systems and hubness. (*a*) A spring-embedded layout of minimum spanning tree for the original network with no distance correction. Each node corresponds to a brain region, and colors indicate the system to which that region is assigned. (*b*) A surface representation of system labels from panel *a*. (*c*) Similar to panel *a* except color now indicates nodes’ ranked participation coefficients calculated using the original network and with respect to the canonically defined brain systems. (*d*) The difference in ranked participation coefficient after correcting for distance. (*e*) Participation coefficient differences projected onto the cortical surface. (*f*) Mean difference in participation coefficient aggregated by system. Opaque and labeled systems are those whose mean difference exceeded chance levels (false discovery rate controlled at 5%).

In typical analyses of functional networks, connections of all lengths are treated equally, making it difficult to parse the unique contributions of long- and short-range connections to any topological feature [4]. Our findings suggest that short-range and long-distance connections make differential contributions to the “hubness” of individual brain regions. Specifically, we find that shifting focus onto long-distance connections results in system-wide increases within participation of primary sensory systems. This observation is counter to the traditional classification of these systems as non-hubs [7, 8], a classification that our findings suggest is likely driven by the strong short-range connectivity among the brain regions that comprise those systems. Notably, our observations are in agreement with tract-tracing studies in animal models that have reported long-distance projections visual and sensorimotor areas [37, 38]. Collectively, our findings suggest an alternative and distance-dependent interpretation of brain areas’ functional roles within the broader context of the network.

### Incorporating distance-dependence into community detection tools

In the previous section, we treated the system labels [36] though they are equivalent to communities. While this approach is useful, communities can also be defined by data-driven methods. Here, we use the community detection heuristic “modularity maximization” to identify and compare communities in the observed and distance-corrected functional connectivity matrices [39].

Specifically, we used a variant of modularity maximization known to work well with correlation matrices [40] and that we have applied to functional connectivity matrices in previous papers [16, 22, 41]. To ensure that comparisons between the two matrices are as appropriate as possible, we guided the modularity maximization algorithm so that it detected a prescribed number of communities, *k* (by selectively varying its resolution parameter, *γ*). This additional constraint guaranteed that in every comparison between the original and distance-corrected networks was made using partitions that resulted in an equal number of communities (Figure, 4a).

Each comparison entailed several steps. First, we computed the pairwise similarity of the partitions using the z-score of the Rand index [42] (see **Materials and Methods**). In general, larger z-scores indicated higher levels of similarity. We used this measure to compare partitions detected using the original and distance-corrected matrices (Figure. 4b). In general, we found that partitions detected using the two techniques were similar.

**FIG. 4.**
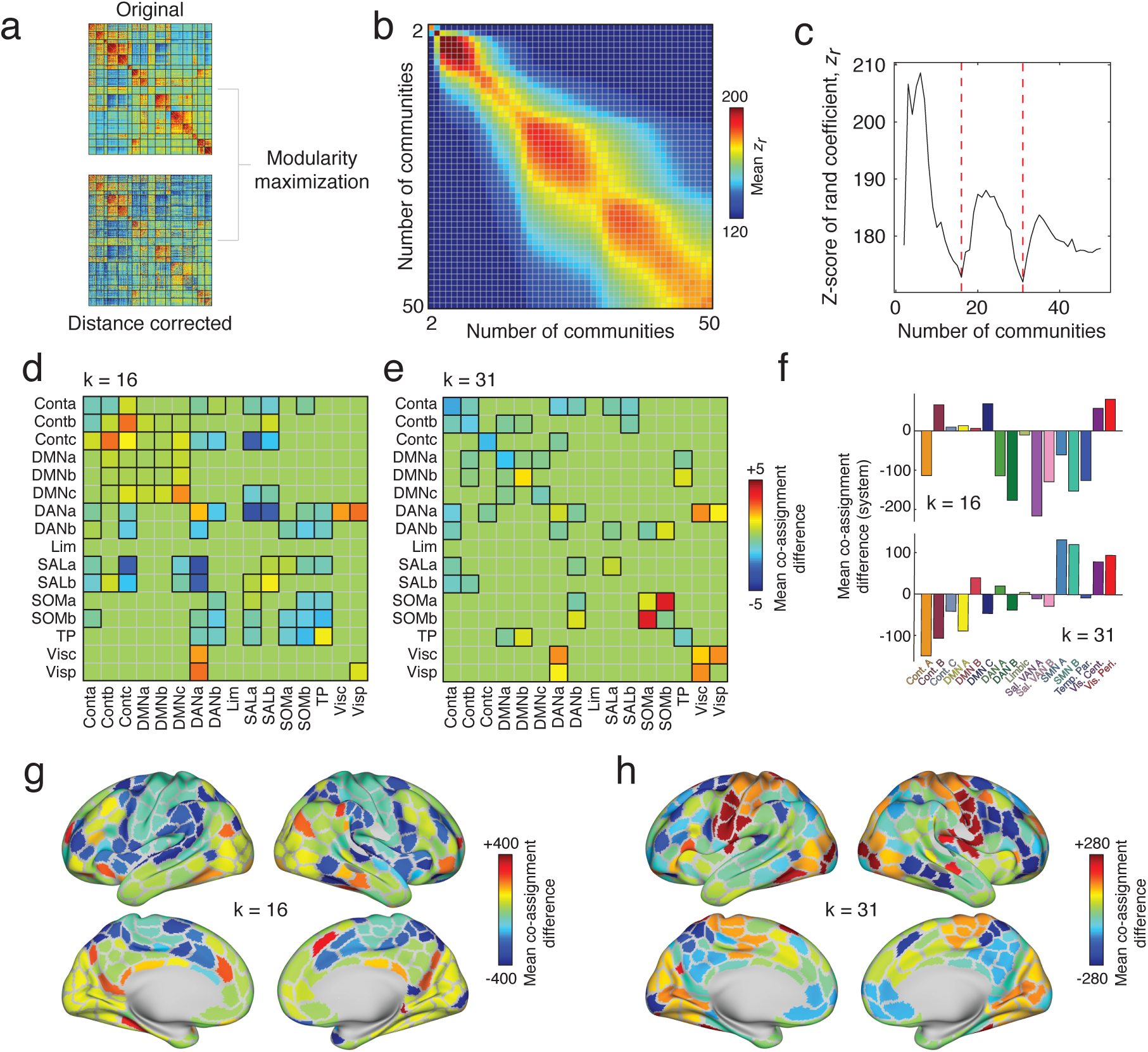
Data-driven estimation and comparison of community structure. (*a*) We used modularity maximization to detect communities in the original and distance-corrected FC matrices. (*b*) We arranged community partitions based on the number of detected communities and compared them using the z-score of the Rand index. Here, we show the mean similarity as a function of the number of communities. (*c*) To narrow our comparisons, we focused on partitions of the brain into 16 and 31 communities, which corresponded to local minima along the diagonal of the matrix shown in panel *b*. Panels *d* and *e* show statistically significant differences in mean module co-assignment. Here and in all subsequent plots, the differences in co-assignment values reflect distance-corrected minus original. Thus, warm colors indicate nodes/systems that are more likely to be co-assigned after correcting for distance; cool colors reflect greater co-assignment in the original matrix than in the distance-corrected matrix. Pairs of systems that survive statistical comparisons have black borders. To better identify those regions and systems whose community co-assignment differed the greatest, we calculated the mean co-assignment difference for every brain region with all other brain regions. In panel *f* we further average those mean differences by system for the *k* = 16 (*top*) and *k* = 31 (*bottom*) partitions. In panels *g* and *h*, we show mean differences plotted on the cortical surface.

Nonetheless, there were subtle differences between detected partitions. To better characterize these differences, we identified the numbers of communities present for which similarity was lowest. We found that these minima occurred when the number of communities were *k* = 16 and *k* = 31 (Figure. 4c). To identify the community features that were driving this dissimilarity, we computed the difference in community co-assignment matrices. Briefly, a co-assignment matrix (alternatively referred to as an “agreement”, “consensus”, or “allegiance” matrix) counts the fraction of times that a pair of nodes were co-assigned to the same community given an ensemble of partitions. For non-deterministic community detection methods like modularity maximization, the co-assignment matrix serves as a pseudo-continuous way of partitioning the network into communities. Comparing co-assignment matrices allows us to identify differences in communities between the two matrices.

To provide better context, we aggregated differences in co-assignment probability by brain systems and compared mean co-assignment differences to those obtained using a permutation-based null model (*p <* 0.05; false discovery rate fixed at 5%) (Fig. 4d,e). In both cases, we observed many subtle yet significant differences. Among the most salient when *k* = 16 were decreases in the co-assignment probabilities of brain areas in the salience network with those in the dorsal attention system and increased co-assignment probability of control network sub-components with one another. In other words, regions associated with these respective networks were more likely to be observed in the same community after correcting for distance than in the original network. Similarly, when *k* = 31, we found increased co-clustering probability within the broader visual and somatomotor systems.

We summarized these results further, calculating the average co-assignment difference for each brain region and subsequently averaging these values for every system (Fig. 4f,g,h). As expected, when *k* = 16, the systems with the greatest co-assignment decreases were concentrated in the salience and dorsal attention systems (Fig. 4f, top), whereas with *k* = 31, the greatest biggest changes occurred within somatomotor and visual networks (Fig. 4f, bottom).

Collectively, these results suggest that differences in observed and distance-corrected functional connectivity are subtle, as evidenced by the high z-score Rand indices. However, these differences are also systematic and emphasized within and between particular brain systems. Moreover, these differences span multiple organizational scales [43]. As in the previous section, these findings suggest that accounting for spatially-driven patterns of FC and focusing on long-distance patterns of inter-areal coupling reveals novel network features; in this case, mani-festing in the form of novel community structure.

### Multi-scale changes in participation coefficient and distinct contributions from somatomotor and visual systems

In the previous two sections, we demonstrated that by comparing observed and spatially constrained patterns of functional connectivity, we were able to tease apart features of networks that are driven differentially by short-range or long-distance connections. Our results also suggested that differences in community structure were manifest at multiple topological scales. This observation is consistent with the idea that brain networks exhibit neuroscientifically relevant features at and across different levels of organization [43]. Here, we pursue this idea further and characterize multi-scale structure of brain areas’ participation coefficients and the network’s hub distribution.

To explore multi-scale hub structure, we first calculated the difference in participation coefficient estimated using the distance-corrected and distance-uncorrected networks. We performed this analysis separately for each brain region and, because we were interested in relative and not absolute changes in participation coefficient, we rank-transformed these values. We then repeated this procedure while varying the number of communities from 2 to 150 (Fig. 5a). Then, we used non-negative matrix factorization (NMF) to generate low-rank approximations of the participation coefficient differences. We compared how well the reconstructed data fit the observed data and found that increasing the number of dimensions beyond four led to little improvement in the overall fit. We repeated the algorithm 100 times and identified, from those repetitions, the group of factors that best reconstructed the original data.

**FIG. 5.**
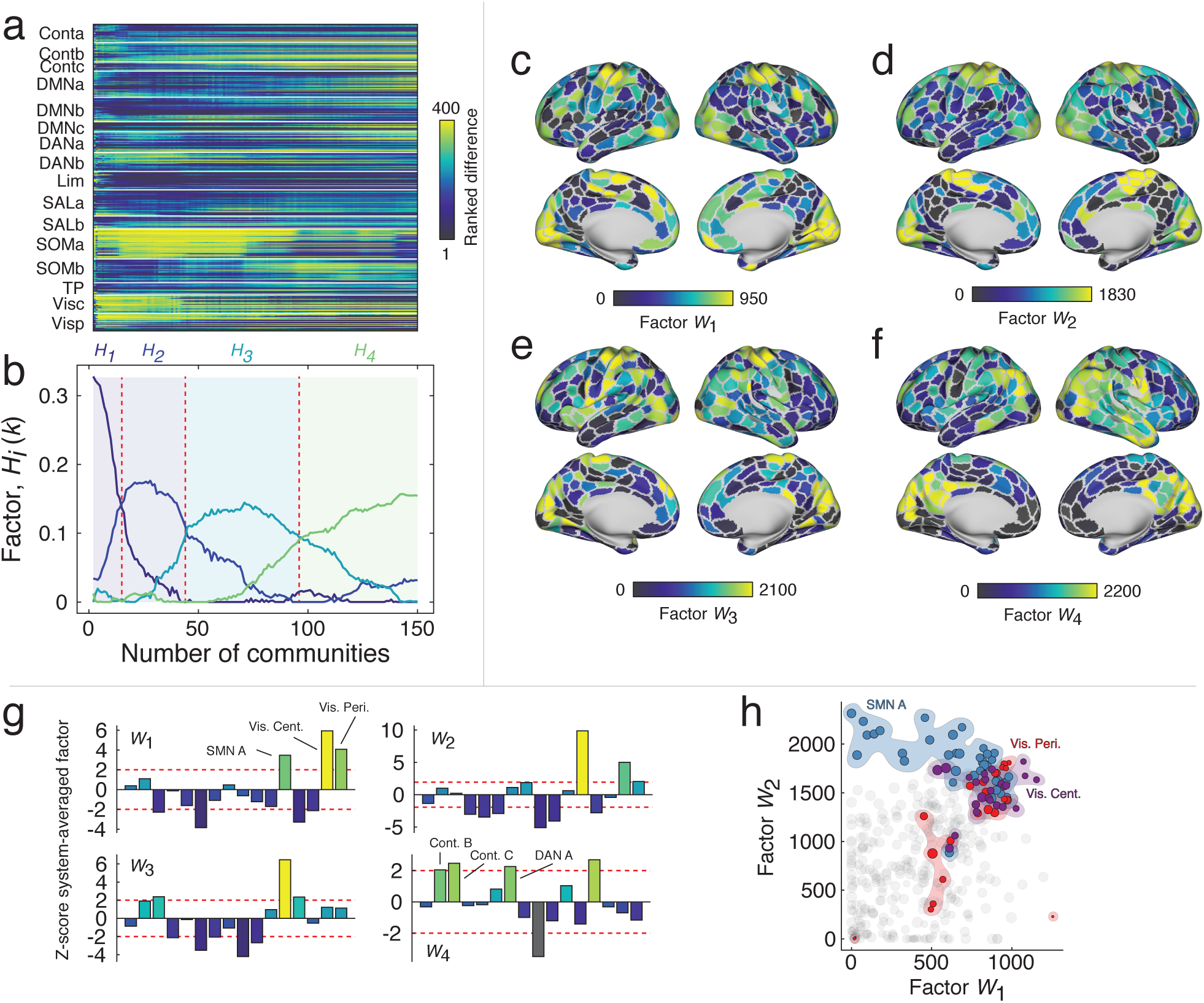
Results of non-negative matrix factorization. (*a*) Ranked differences in participation coefficients for all brain regions and with the number of communities ranging from 2 to 150. This matrix served as input to the NMF algorithm. (*b*) NMF identified four factors that approximately reconstructed the original data. Broadly, these factors were associated with different topological regimes corresponding to different numbers of communities. In panels *c*-*f* we show the factors mapped onto the cortical surface. Note that the first three factors emphasize participation coefficient differences in motor and visual systems while the fourth factor highlights widespread, multi-system differences. (*g*) We show system-averaged and z-scored differences in participation coefficient. Note that the first three factors all emphasize motor and visual systems but in different proportions. (*h*) Scatterplot of first two factors against one another. Note that somatomotor and visual areas tend to display the largest values in general. However, note also that those areas deviate from the diagonal, indicating that differences in these systems are emphasized differentially by the two factors and at different topological scales.

Analyzing those four factors further, we found that their expression varied more or less smoothly as a function of the number of communities (Fig. 5b). Broadly speaking, the first three factors were similar to one another, in that they emphasized increased participation coefficient within visual and somatomotor areas. However, the extent to which those systems and respective subsystems was expressed varied (Fig. 5c-e). For instance, the first factor (*W*_1_) exhibited differences in the participation coefficient of visual areas and peaked earlier, in terms community number, than factors *W*_2_ and *W*_3_ (Fig. 5g). Those factors, on the other hand, peaked later and emphasized the differences in somatomotor participation coefficient. These differences are made more pointed when we directly compare *W*_1_ with *W*_2_ in a scatterplot (Fig. 5h). We find, in both cases, that visual and motor areas are among the strongest values, but that the somatomotor system (SMN A) falls to the left of the main diagonal (stronger in *W*_2_) while the visual system (Vis-Cent) falls to the right of the main diagonal (stronger in *W*_1_). The fourth factor, on the other hand, peaked the latest and exhibited neither motor nor visual systems, but more spatially diffuse differences among control and dorsal attention systems (Fig. 5f,g).

The import of these findings spans several domains. First, the results suggest that space-corrected hub structure varies, as a function of the number of detected communities., albeit subtly. This observation agrees with recent multi-scale accounts of brain network structure that have suggested that the brain exhibits unique and functionally relevant organizational features at multiple topological scales [43]. Multi-scale and hierarchical organization is critical for complex systems, as it engenders robustness to perturbations and separation of dynamical timescales [44], properties that are essential for embodied nervous systems that interact with variable environments.

## DISCUSSION

In this report, we analyzed the contributions of long-distance connectivity to the community and hub structure of functional brain networks. To do this, we developed a surrogate-based method that generates synthetic networks lacking long-distance correlations but preserving short-range connectivity. We found that even these null networks exhibited neuroscientifically interesting structure, including strong correlations within primary sensory systems, such as visual and somatomotor areas, as well as sub-components of higher-order cognitive systems, such control networks. These observations suggested that connections of different lengths contribute to shape the overall character of the network.

Next, to better understand those distinct contributions, we suppressed the contributions from short-range connections by constructing a “distance-corrected” connectivity matrix in which we simply subtracted the short-range synthetic network from the observed. The resulting network expressed long-distance connections but effectively suppressed those made between proximal brain areas. We compared the features of the distance-corrected network to those of the observed network and found that the participation coefficient of visual and somato-motor regions increased in the distance-corrected network, suggesting that the long-distance connectivity patterns of these brain areas leave those systems increasingly well-situated for integrating information across communities. We also found evidence that the community structures of the two networks differed across multiple topological scales, confirming further that long-distance and short-range connections uniquely shape the organization of functional networks. In summary, our work shows that the weights of functional connections exhibit a complicated relationship with space and distance, but also presents a flexible corrective strategy for teasing apart the contributions of spatial sources to functional connectivity. Our approach outlines a procedure that enables future studies to conduct a more nuanced analysis of functional networks in both health and disease.

### Spatial constraints and brain connectivity

Here, we present a careful analysis of functional networks in which we discount the effect of short-range connections. Why bother doing so? What can we learn about brain organization and function by studying connections of different lengths? The primary justification for focusing on longer connections is that they entail greater costs than short connections (all other things equal) and that their mere existence suggests functional relevance [17, 45]. This is certainly true in the case of structural and anatomical connections, where the metabolic and material costs of forming and maintaining axonal projections or fiber tracts grow as a function of their length and diameter [18, 46, 47]. It is also likely true in the case of functional connections, which are the focus of this study and which require energy to maintain correlated activity over long distances, especially in the absence of direct structural support [48]. In this light, the analyses presented can be viewed as effectively shifting focus onto the more costly features of functional networks and ones that, we argue, are of greater relevance to the overall network function.

We acknowledge that in focusing on long and costly functional connections, we necessarily overlook network features that are driven by short-range connectivity. This is not to say that short connections are irrelevant to network function. Short-range connections are generally among the strongest in functional networks, and they factor disproportionately in the weighted shortest path structure of anatomical networks [16], and enhancing the cohesiveness of communities [10]. Accordingly, we view short-range and long-distance connections as complementary, with each delivering unique insights into the organization and behavior of functional networks.

In addition to characterizing the organization of long-distance functional connections, our study also contributes some useful methodology. An important component of any network neuroscience study is the comparison of some measurement made on the observed network with a null distribution of the same measure made on an ensemble of appropriately constructed random networks. Because the randomized networks tend to preserve only low-level features of the observed network, e.g. average binary density, degree sequence, etc., this hypothesis testing framework allows us to identify higher-level features of the observed network that are not easily attributable to random fluctuations [49]. Increasingly, it is becoming understood that the traditional random network models can be too liberal for many of the hypothesis testings [50–53]. That is, though they preserve degree (in un-weighted network) or strength (in weighted network) sequences, they also fail to preserve other key attributes of real-world brain networks, including wiring cost, or violate mathematical relationships when applied to correlation matrices [33]. These failures can result in mischaracterizations of networks, which identify features that act as “spandrels” and emerge as a result of relatively benign processes. The spatial null model we use here addresses some of these concerns, as it generates admissible correlation matrices and allows the user to flexibly model spatial relationships. In future work, similar null models could be used both to identify novel network features and better clarify the relationship of known features with psychological phenomena and brain function.

### Increased participation coefficient in primary sensory regions

One of the most salient findings we report is the increased participation coefficient of brain regions comprising the visual and somatomotor systems. The conventional interpretation of high-participation nodes is that they serve as network “hubs” [7, 54]; their connections form bridges across multiple sub-systems, which engenders or reflects the polyfunctionality of those regions [9, 35]. In past studies, somatomotor and visual areas were generally reported among the least hub-like parts of the brain; regions in those systems made strong connections within their respective communities but weak extra-community connections [7, 8, 54]. Our findings, on the other hand, suggest that the low levels of “hubness” in those regions is a direct consequence of their strong short-range connectivity and that, in terms of their longer connections, visual and motor systems can be regarded as much more hub-like. These observations suggest expanded functional roles for these systems, which are traditionally associated with uni-modal information processing. Future work should investigate this question more directly.

### A less severe approach to control for spatial artifacts

Occasionally spatial relationships have been discussed in the context of FC, most often with respect to the origins of short-range FC. Many influential studies, for instance, regard short-range FC as having artifactual origins and, as a way of addressing such artifacts, simply discard all FC between regions separated by distances less than some threshold, e.g. 20 millimeters [6]. While this strategy is effective in reducing the number of false-positives (artifactually strong short-range correlations), it may inadvertently discard true-positives and does so in a binary way – connections are either retained or discarded, with no middle ground. The strategy we propose here, on the other hand, offers a graded and more systematic way of taking into account the spatial proximity of connections in functional connectivity analysis. Specifically, the severity with which we discount a connection’s observed weight decays monotonically with the distance between nodes. So the closer two nodes are to one another, the greater the extent to which they are discounted. In this sense, our strategy may prove more beneficial, in that the continuous discounting of connections does not throw away connectivity information; the short-range connections can still contribute to estimates of community structure or participation, but with less overall influence.

### Limitations

Though this study makes several important contributions, it also has a number of limitations. First, this study focuses on group-averaged FC rather than FC of single subjects. While this strategy facilitates computation (we only have to fit the models to a single connectivity matrix), the lack of single-subject analyses means that we may fail to appreciate the individual variability in weight-distance dependencies [55] and also run the risk of characterizing patterns in the group matrix that are not representative of any typical subject [56]. Indeed, future applied studies are needed to determine how results reported here map onto individual subjects and how those mappings are related to inter-individual variability in subjects’ performances on psychometric and cognitive tests.

Another challenge involves processing decisions made as part of modeling distance-dependent FC. Essentially, we treat the observed FC as though it were made up of three components: “extra-spatial” + “spatial” + “error” FC. Our aim was, as best as possible, to subtract away the spatial component, leaving only the true FC and error terms. The challenge, however, is that we do not know the actual forms of *any* of the three components nor do we know the function by which they are intermixed. Here, we assume that the spatially-driven FC decays exponentially with distance and that by 60 millimeters its effect is small [31]. We also assume that the spatial component enters linearly, so that its effect can be subtracted out. While these assumptions were made with practical considerations in mind, such as model parsimony and computational complexity, future work should investigate and test these assumptions systematically and explicitly.

Finally, here we have used Euclidean distance to obtain the weighted contribution of different brain areas. However, previous studies [57] suggest that using Euclidean distance could potentially reduce the spatial selectivity meaning that functionally different regions could be selected by voxel selection methods that depend on Euclidean distance. Therefore, other distance measures such as geodesic distance that respect the curvature of the cortex during calculations should be studied and its performance compared to Euclidean distance.

## MATERIALS AND METHODS

### Human Connectome Dataset

In this study, we aimed to characterize the relationship between space and resting state FC. To address this aim, we leveraged data from the Human Connectome Project (HCP), a multi-site consortia that collected extensive MRI, behavioral, and demographic data from a large cohort of subjects (*>*1000) [58]. As part of the HCP protocol, subjects underwent two separate resting state scans along with seven task fMRI scans. All functional connectivity data analyzed in this report came from these scans and was part of the HCP S1200 release [58]. Subjects that completed both resting-state scans and all task scans were analyzed. We utilized a cortical parcellation that maximizes the similarity of functional connectivity within each parcel (*N* = 400 parcels) [36].

We processed ICA-FIX data provided by the HCP, which used ICA to remove nuisance and motion signals [59]. In addition, the 12 detrended motion estimates provided by the Human Connectome Project were regressed out from the time series, the mean global signal was removed, and the time series was bandpass filtered from 0.009 to 0.08 Hz.

For all scans, the MSMAII registration was used, and the mean time series of vertices on the cortical surface (fsL32K) in each parcel was calculated. These time series were z-scored and concatenated across all subjects. A group representative functional connectivity matrix was then calculated as the pairwise Pearson correlation (subsequently Fisher *z*-transformed) between concatenated node time series. We denote this matrix as *A* and whose element *A*_*ij*_ refers to the Fisher-transformed correlation between regions *i* and *j*.

### Modeling distance dependence

Our goal was to understand the role of spatial relationships in shaping whole-brain FC. To investigate this question, we constructed a distance-based null model. We reasoned that this model should have the following properties: 1) FC should decay monotonically with distance, so that proximal brain areas are more strongly correlated than distant brain areas; 2) the connectivity matrix must be admissible as a correlation matrix [33]. These requirements are in contrast to current null models for FC, in which binarized connections are rewired while preserving degree – an operation that can result in edge configurations that violate transitive relationships associated with the correlation metric [31].

There are many ways of realizing such a model, though here we focused on a single possibility based on surrogate time series analysis. Suppose *x*_*i*_ = [*x*_*i*_(*t*)] is the observed activity recorded from brain area *i ∈ N*. We could generate a surrogate time series with the same power spectrum by taking the discrete Fourier transform of *x*_*i*_, adding or subtracting random amounts of phase to each frequency bin, and performing an inverse Fourier transform. We denote the resulting time series as 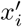. We note that the phase-randomization surrogate procedure was always carried out at the level of individual subjects; all surrogate time series were subsequently concatenated.

If we performed this operation independently and for each brain area, the resulting inter-areal correlations would be weak, and the full matrix would exhibit no structure. To induce spatial dependencies, we generate for each brain area another time series, 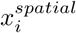, that is the linear combination of every other regions’ time series, but where the mixing coefficients are distance-dependent:

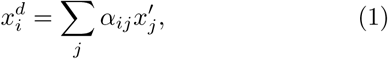

where *α*_*ij*_ is defined according to the function *f* (*D*_*ij*_, *β*). Here, *D*_*ij*_ is the Euclidean distance between nodes *i* and *j*, and *β* is the parameter that controls the decay rate of FC with distance. In general, we define this function in any way, but for practical reasons and to maintain consistency with previous work, we defined it to be the exponential: *f* (*D*_*ij*_, *β*) = exp(−*β · D*_*ij*_) [30, 60].

Fitting this model amounted to choosing the *β* parameter that optimizes some objective function. Here, we used the bisection method to maximize the correlation of the observed FC matrix, *A*, with the distance-dependent FC matrix, *A*^*spatial*^. We defined model fitness as the average fitness over 50 independently generated surrogate time series (optimal fitness of *r* = 0.35 at *β* = 0.078).

### Modularity Maximization

It is generally understood that brain networks can be decomposed into clusters defined based on nodes’ connectivity patterns. These clusters, also called communities or modules, are typically unknown ahead of time and estimated using data-driven approaches. Of so-called community detection methods, modularity maximization remains one of the most widely used. Modularity maximization operates on a simple principle: compare the connectivity patterns we observe with those we would expect by chance. Communities are defined as groups of nodes more strongly connected to one another than expected. This intuition can be formalized by the modularity heuristic. If *B*_*ij*_ = *A*_*ij*_ − *P*_*ij*_ represents the difference in weight of the observed and expected connection between node *i* and *j*, then we can define the following quality function:

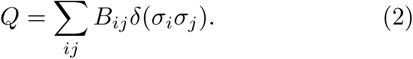

In this expression, *δ*(*·, ·*) is the Kronecker delta function, whose value is 1 if its arguments are equal and 0 otherwise. Here, its arguments are the *σ*_*i*_ and *σ*_*j*_, which denote the community assignments of nodes *i* and *j*. As a result of the delta function, the only elements *B*_*ij*_ that contribute to the summation are those that fall within communities.

The value of *Q* can be used to rate the quality of a proposed community partition. Alternatively, an optimal community structure can be uncovered by assigning nodes to the communities that optimize *Q*. This optimization procedure is computationally intractable, though the optimal solution can be approximated using several heuristics, the most popular of which is the so-called “Louvain algorithm”: a non-deterministic and agglomerative method that generally performs well on most benchmarking tests.

Here, we use a modularity framework that has been extended in several important ways. First, we focus on a “multi-scale” version in which a structural resolution parameter, *γ*, is introduced to the modularity matrix: *B*_*ij*_ = *A*_*ij*_ − *γP*_*ij*_. The value of *γ* can be tuned to smaller or larger values to, effectively, uncover communities of correspondingly larger or smaller size, respectively. Second, we define the expected weight of connections to be *P*_*ij*_ = 1 for all {*i, j*}. This particular null model has proven to be compatible with correlation matrices and results in an intuitive definition of communities as groups of nodes whose average internal density exceeds a value of *γ*. To detect the community structure of observed FC (uncorrected for distance), we optimized the following multi-scale modularity index:

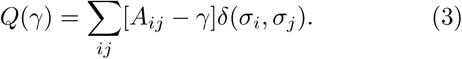

#### Distance-dependent modularity maximization

We optimized a similar modularity index to detect communities for distance-corrected FC by defining the modularity matrix: 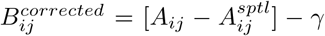. The distance-corrected modularity can then be expressed as:

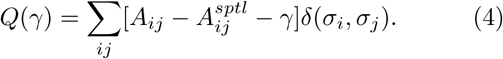

We optimized both the original and distance-dependent modularity using a generalized version of the Louvain algorithm. Typically, as part of multi-scale community detection, the value of *γ* is either systematically or randomly sampled over some predefined range. Here, we aimed to compare the multi-scale community structure of distance-corrected and uncorrected FC. To ensure that the comparison was as fair as possible, e.g., not comparing a partition of the network into two communities with another that divides the network into twenty communities, we used an adaptive algorithm to sample partitions of both distance-corrected and uncorrected FC into the same number of communities.

To do so, we used a two-step adaptive algorithm. First, we coarsely sampled 1001 different *γ* values over the range of *γ* = *−*0.2 to *γ* = 0.7, which encompasses the range of neuroscientifically interesting partitions. We optimized *Q*(*γ*) for each network and for each value of *γ* and calculated the number of communities in the resulting partition. Using this approach, we obtained a rough mapping of *γ* to the number of communities. This mapping allowed us to select a number of communities, e.g., *k* = 14, and specify a range of *γ* values for which we could reasonably expect to obtain 14 communities. Finally, we varied the value of *k* from 2 to 150, sampling *γ* from restricted ranges until, for each value of *k*, we sampled 250. We repeated this procedure for both the uncorrected and distance-corrected FC matrices, resulting in two sets of 37,500 partitions in total.

#### Z-scored Rand index for partition similarity

We used the z-score of the Rand index to measure the similarity of partitions detected using the uncorrected and distance-corrected FC matrices. This measure is similar to the traditional Rand index but corrects for biases induced by the number and size of the communities in the partitions being compared. For two partitions, *X* and *Y*, we measure their similarity as:

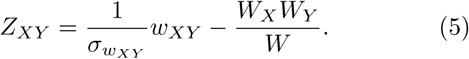

Here, *W* is the total number of node pairs in the network, *W*_*X*_ and *W*_*Y*_ are the number of pairs in the same modules in partitions *X* and *Y*, respectively, *w*_*XY*_ is the number of pairs assigned to the same module in *both X* and *Y*, and 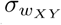 is the standard deviation of *w*_*XY*_. The value of *Z*_*XY*_ can be interpreted as how great, beyond chance, is the similarity of partitions *X* and *Y*.

#### Participation coefficient

Using the detected communities and based on FC, we can also identify those brain regions whose connections span the boundaries of communities (polyfunctional) and those whose connections are confined, largely, to their own community (unifunctional). To identify these kinds of brain regions, we calculated the network measure *participation coefficient* for each brain region, *i*, which we denote as *P*_*i*_:

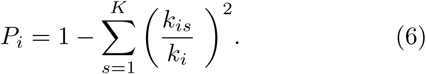

Here, *k*_*i*_ = Σ_*j*_ *A*_*ij*_ is node *i*’s weighted degree and *k*_*is*_ = Σ_*j∈s*_ *A*_*ij*_ is the total weight of node *i*’s connections to module *s*. Participation coefficients range from 0 to 1, where larger values indicate that connections are evenly spread over modules. Here, because our matrices were signed, we used a modified version of participation coefficient that considers only positive connections. To ensure that comparisons were unbiased by the effect of weight, we rank-transformed brain regions’ participation coefficients.

### Non-negative matrix factorization

Non-negative matrix factorization (NMF) is a technique that generates low-rank approximations of a, potentially, high-dimensional dataset, **X** ∈ ℝ^*n×p*^. Briefly, this approach entails identifying matrices **W** ∈ ℝ^*n×k*^ and **H** *∈* ℝ^*k×p*^ such that **W***×***H** *≈* **X** and subject to the constraint that all elements of **W** and **H** are non-negative. Here, we used NMF to decompose brain-wide participation coefficients as the number of communities varied from 2 to 150. Thus the dataset had dimensions **X** ∈ ℝ^400×149^. The MATLAB implementation of NMF uses the non-deterministic alternating least squares algorithm to determine **W** and **H**. Consequently, we repeated the algorithm 100 times with different initial conditions as we varied the rank from *k* = 2 to *k* = 20. We observed that the root mean square of the residual (a measure of fitness) decreased sharply until *k* = 4, suggesting that the original dataset could be reasonably approximated using four dimensions.

## ACKNOWLEDGMENTS

This research was supported by Indiana University Office of the Vice President for Research Emerging Area of Research Initiative, Learning: Brains, Machines and Children (FZE and RFB). DSB acknowledges support from the John D. and Catherine T. MacArthur Foundation, the Alfred P. Sloan Foundation, the Paul Allen Foundation, the Army Research Laboratory through contract number W911NF-10-2-0022, the Army Research Office through contract numbers W911NF-14-1-0679 and W911NF-16-1-0474, the National Institute of Health (2-R01-DC-009209-11, 1R01HD086888-01, R01-MH107235, R01-MH107703, R01MH109520, 1R01NS099348 and R21-M MH-106799), the Office of Naval Research, and the National Science Foundation (BCS-1441502, CAREER PHY-1554488, BCS-1631550, and CNS-1626008). The content is solely the responsibility of the authors and does not necessarily represent the official views of any of the funding agencies.

